## Supplemental data for "Nanotiming: telomere-to-telomere DNA replication timing profiling by nanopore sequencing"

#### Supplemental information

**Supplementary Figure 1. Comparison between mean BrdU content and sort-seq relative copy number profiles of all yeast chromosomes.** See Fig. 2 caption for details. BT1 wt_rep1 mean BrdU content profile is shown.

**Supplementary Figure 2. Comparison between mean BrdU content and MFA-seq relative copy number profiles of all yeast chromosomes.** See Fig. 2 caption for details. BT1 wt_rep1 mean BrdU content profile is shown.

**Supplementary Figure 3. Reproducibility of mean BrdU content profiling of BT1 chromosomes.** Mean BrdU content was computed in 1 kb bins along nanopore reads of genomic DNA of six independent BT1 cell cultures labelled for one doubling time with 5 µM BrdU. See Fig. 2 caption for details.

**Supplementary Figure 4. Spearman's rank correlation coefficients of pairwise comparisons between six independent mean BrdU content profiles, one relative copy number profile from sort-seq and one relative copy number profile from MFA-seq of *S. cerevisiae* genome.** Mean BrdU content and sort-seq profiles were computed from reads of genomic DNA from BT1 cells; MFA-seq data are from ref.[8]. NanoT, Nanotiming; wt, wild-type; rep, replicate.

**Supplementary Figure 5. Mean BrdU content profiles of all chromosomes of wild-type and *ctf19*Δ BT1 cells.** See Fig. 2 caption for details.

**Supplementary Figure 6. Mean BrdU content profiles of all chromosomes of wild-type and *rif1*Δ BT1 cells.** See Fig. 2 caption for details.

**Supplementary Figure 7. Mean BrdU content profiles of all chromosomes of wild-type and *yku70*Δ BT1 cells.** See Fig. 2 caption for details.

**Supplementary Figure 8. Mean BrdU content profiles of all chromosomes of wild-type and *fkh1*Δ BT1 cells.** See Fig. 2 caption for details.

**Supplementary Figure 9. Comparison between mean BrdU content and sort-seq relative copy number profiles of all chromosomes of *S. cerevisiae ctf19*Δ cells.** Mean BrdU content profile was computed from reads of genomic DNA from *ctf19*Δ BT1 cells (rep3); sort-seq data are from ref.[21]. See Fig. 2 caption for details.

**Supplementary Figure 10. Comparison between mean BrdU content and sort-seq relative copy number profiles of all chromosomes of *S. cerevisiae rif1*Δ cells.** Mean BrdU content profile was computed from reads of genomic DNA from *rif1*Δ BT1 cells (rep1); sort-seq data are from ref.[7]. See Fig. 2 caption for details.

**Supplementary Figure 11. Spearman's rank correlation coefficients of pairwise comparisons between three independent mean BrdU content profiles and one relative copy number profile by sort-seq of *ctf19*Δ and *rif1*Δ genomes. a**, **b**, Results for the genome of *ctf19*Δ (**a**) and *rif1*Δ (**b**) strains. Mean BrdU content profiles were computed from reads of genomic DNA from *ctf19*Δ (**a**) and *rif1*Δ (**b**) BT1 cells; *ctf19*Δ and *rif1*Δ sort-seq data are from ref.[8] and ref.[7], respectively. NanoT, Nanotiming; rep, replicate.

**Supplementary Figure 12. Impact of the amount of sequencing data on Nanotiming RT profile accuracy. a**, **b**, Evolution of Spearman's rank correlation coefficient between mean BrdU content and sort-seq profiles of BT1 genome in wild-type cells as a function either of the number of nanopore reads used to compute mean BrdU content profiles (**a**) or of genomic coverage (**b**). Horizontal dashed line corresponds to a Spearman's rank correlation coefficient of 0.9. Reads were randomly selected from BT1 wt_rep1 dataset; subsampling was performed 100 times for each read number or genomic coverage category. Horizontal black line, median; boxes, 25th to 75th percentiles; whiskers, 1.5x interquartile range. x, fold.

**Supplementary Figure 13. Evaluation of Nanotiming noise as a function of sequencing depth.** Noise estimator corresponds to the variance of signal differences between consecutive 1 kb bins. Reads were randomly selected from the complete BT1 PromethION dataset; subsampling was performed 10 times for each genomic coverage category. To remove noise dependency in successive bins of nanopore reads, which typically span 10 to 20 kb, bin-to-bin signal variation was compared between consecutive bins from independent samplings. Horizontal black line, median; boxes, 25th to 75th percentiles; whiskers, 1.5x interquartile range. Orange and blue dots, noise estimation for MFA-seq and sort-seq profiles, respectively.

MFA-seq data are from ref.[8]. At equivalent genomic coverage, noise level in a Nanotiming profile at 1 kb resolution is 3.3 and over 30 times lower than in a sort-seq and MFA-seq profile, respectively. x, fold. See Methods for details.

**Supplementary Figure 14. Mean BrdU content profiles of all chromosomes of BT1 cells from a multiplexed PromethION run with 24 barcoded samples of BT1 BrdU-labelled DNA.** The corresponding sort-seq relative copy number profile is also shown. See Fig. 2 caption for details.

**Supplementary Figure 15. Spearman's rank correlation coefficients of pairwise comparisons between mean BrdU content profiles of BT1 genome from a multiplexed PromethION run with 24 barcoded samples originating from the same BT1 BrdU-labelled genomic DNA.** Comparison with BT1 genome sort-seq relative copy number profile is also provided. NanoT, Nanotiming; rep, replicate.

**Supplementary Figure 16. Mean BrdU content profiles over 50 kb of the left and right extremities of *S. cerevisiae* chromosomes in wild-type and *rif1*Δ BT1 cells.** See Fig. 3 caption for details.

**Supplementary Figure 17. Yku70 regulates the RT of *S. cerevisiae* X and XY' telomeres.** **a**, RT at individual telomeres in wild-type and *yku70*Δ BT1 cells. **b**, RT at X and XY' telomeres in wild-type and *yku70*Δ cells. See Fig. 3 caption for details. **a**, **b**, TEL13R RT was not determined in *yku70*Δ mutant because of a missing Y' element at the right end of chromosome XIII compared to BT1 assembly, preventing proper read mapping and telomeric RT computation.

**Supplementary Figure 18. Mean BrdU content profiles over 50 kb of the left and right extremities of *S. cerevisiae* chromosomes in wild-type and *yku70*Δ BT1 cells.** See Fig. 3 caption for details. Please note the missing distal Y' element at the right end of chromosome XIII in *yku70*Δ mutant.

**Supplementary Figure 19. Analysis of the relationship between telomere length and RT at the single-telomere level in wild-type BT1 cells.** See Fig. 4 caption for details.

**Supplementary Figure 20. Analysis of the relationship between telomere length and RT at the single-telomere level in *rif1*Δ BT1 cells.** See Fig. 4 caption for details.

**Supplementary Figure 21. Analysis of the relationship between telomere length and RT at the single-telomere level in *yku70*Δ BT1 cells.** See Fig. 4 caption for details. No data was computed at TEL13R because of a missing Y' element at the right end of chromosome XIII compared to BT1 assembly.

**Supplementary Figure 22. Analysis of the relationship between telomere length and RT at the single-telomere level in *ctf19*Δ BT1 cells.** See Fig. 4 caption for details.

**Supplementary Figure 23. Analysis of the relationship between telomere length and RT at the single-telomere level in *fkh1*Δ BT1 cells.** See Fig. 4 caption for details.

**Supplementary Figure 24. Flow cytometry analysis.** **a**, Gating strategy. Cells are fixed in ethanol and DNA is counterstained with SYTOX Green prior to flow cytometry analysis. Cells are initially gated using the FSC-Area versus SSC-Area plot to remove debris (left panel), then interrogated by the ratios of area (Sytox green FITC-A) to height (Sytox green FITC-H) of the SYTOX Green signal to gate out cell doublets (middle panel). A SYTOX Green area (Sytox green FITC-A) histogram shows DNA content after gating (right panel). **b**, Sorting of BT1 cells. Cells in S and G2 phases of the cell cycle, as well as an "All cells" control, were sorted using the indicated gates positioned on the DNA content histogram (left panel). Sorted cell populations are presented on the right panel.

**Supplementary Table 1. Nanopore sequencing costs.** Prices for R9.4.1 chemistry are those we paid in 2023/early 2024 (original prices in euros were converted into US dollars assuming an exchange rate of 1 USD/EUR); estimated library and barcodes prices for R10.4.1 chemistry are from https://store.nanoporetech.com/native-barcoding-kit-96-v14.html (as of July 2024).

**Supplementary Table 2. Spearman's rank correlation coefficients of comparisons between telomere length and RT in wild-type, *rif1*Δ, *yku70*Δ, *ctf19*Δ and *fkh1*Δ BT1 cells.** No data was computed at TEL13R in *yku70*Δ mutant because of a missing Y' element at the right end of chromosome XIII compared to BT1 assembly, preventing proper read mapping. Statistical significance was set to $p < 0.01$. n, number of measurements; rho, Spearman's rank correlation coefficient; RT, replication timing (mean BrdU content data were rescaled between 1 and 2 corresponding to the end and start of S phase, respectively); wt, wild-type.

**Supplementary Table 3. Spearman's rank correlation coefficients of comparisons between telomere length and RT according to their X/XY' status in wild-type, *rif1*Δ, *yku70*Δ, *ctf19*Δ and *fkh1*Δ BT1 cells.** See Supplementary Table 2 caption for details.

**Supplementary Table 4. Spearman's rank correlation coefficients of comparisons between telomere length and RT for each chromosome end in wild-type, *rif1*Δ, *yku70*Δ, *ctf19*Δ and *fkh1*Δ BT1 cells.** See Supplementary Table 2 caption for details.

Supplementary Figure 1

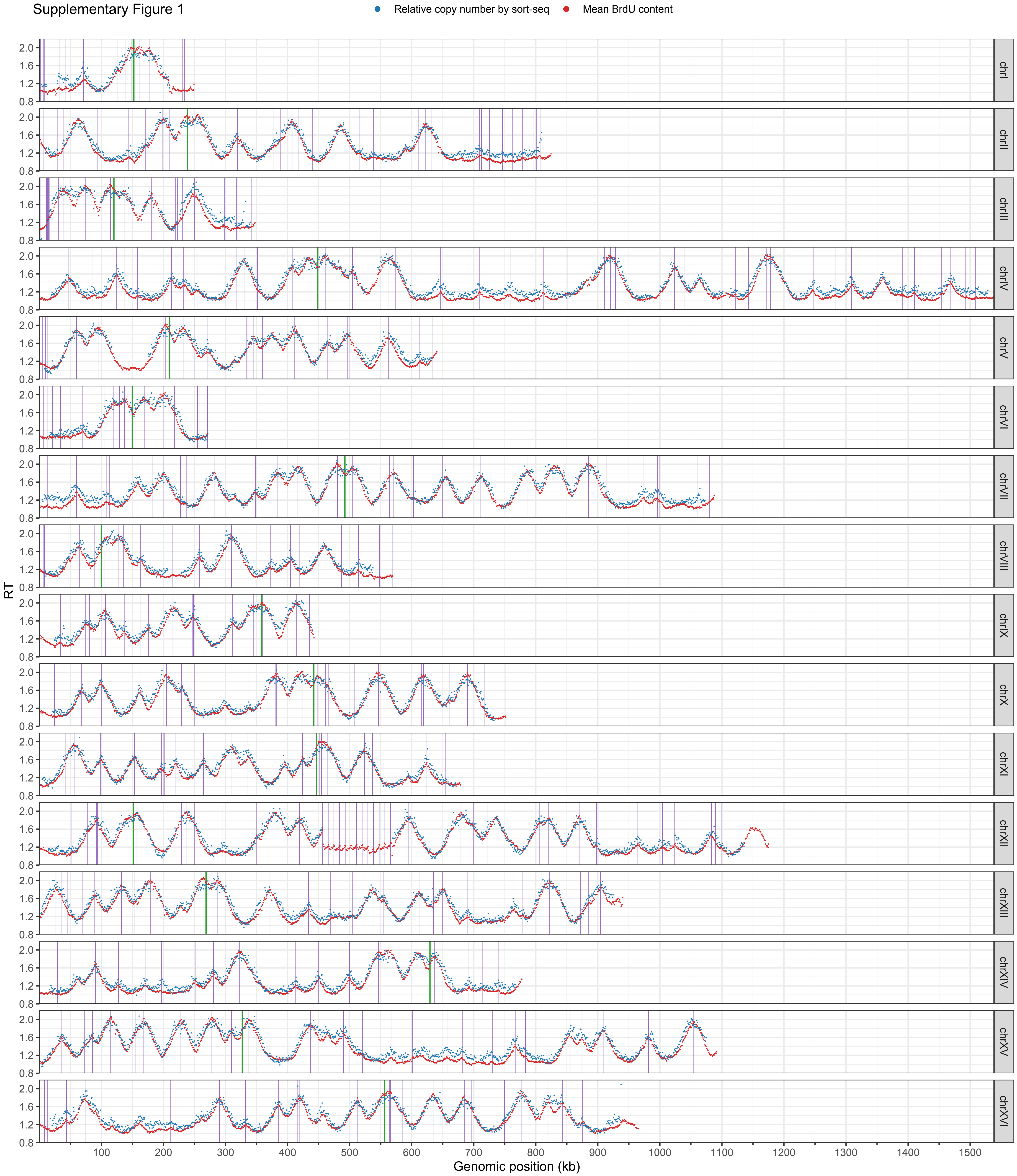

Supplementary Figure 2

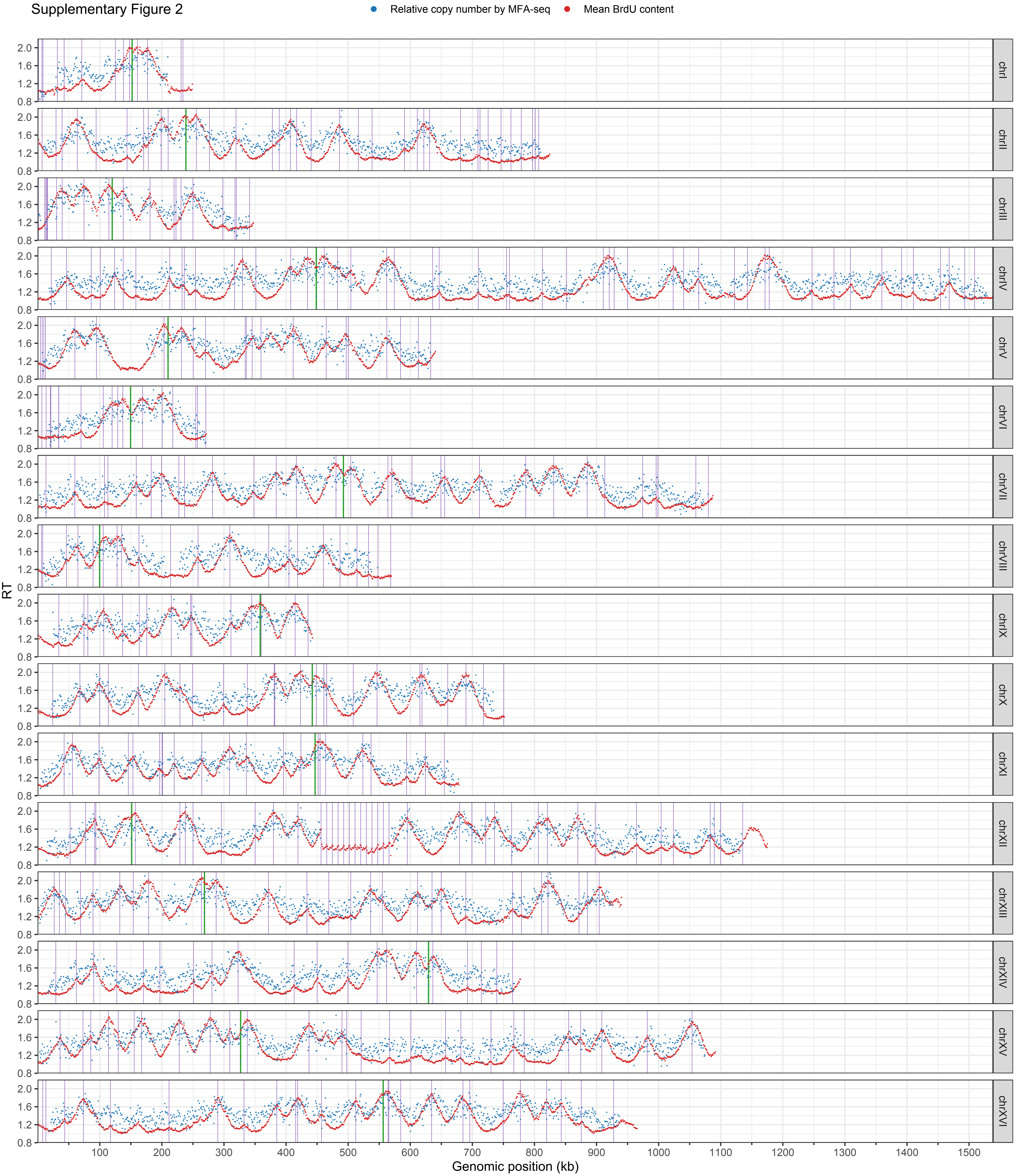

Supplementary Figure 3

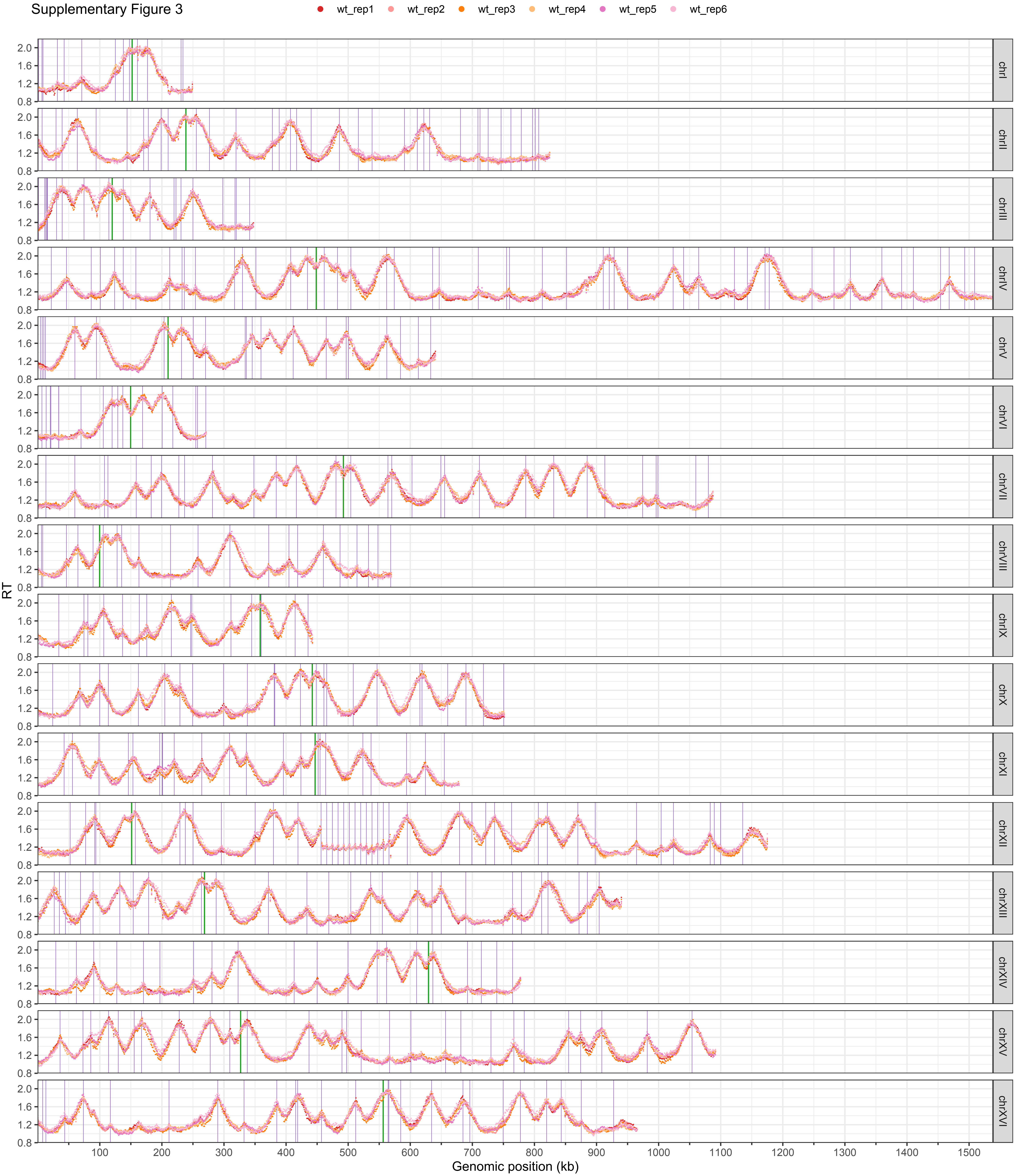

Supplementary Figure 4

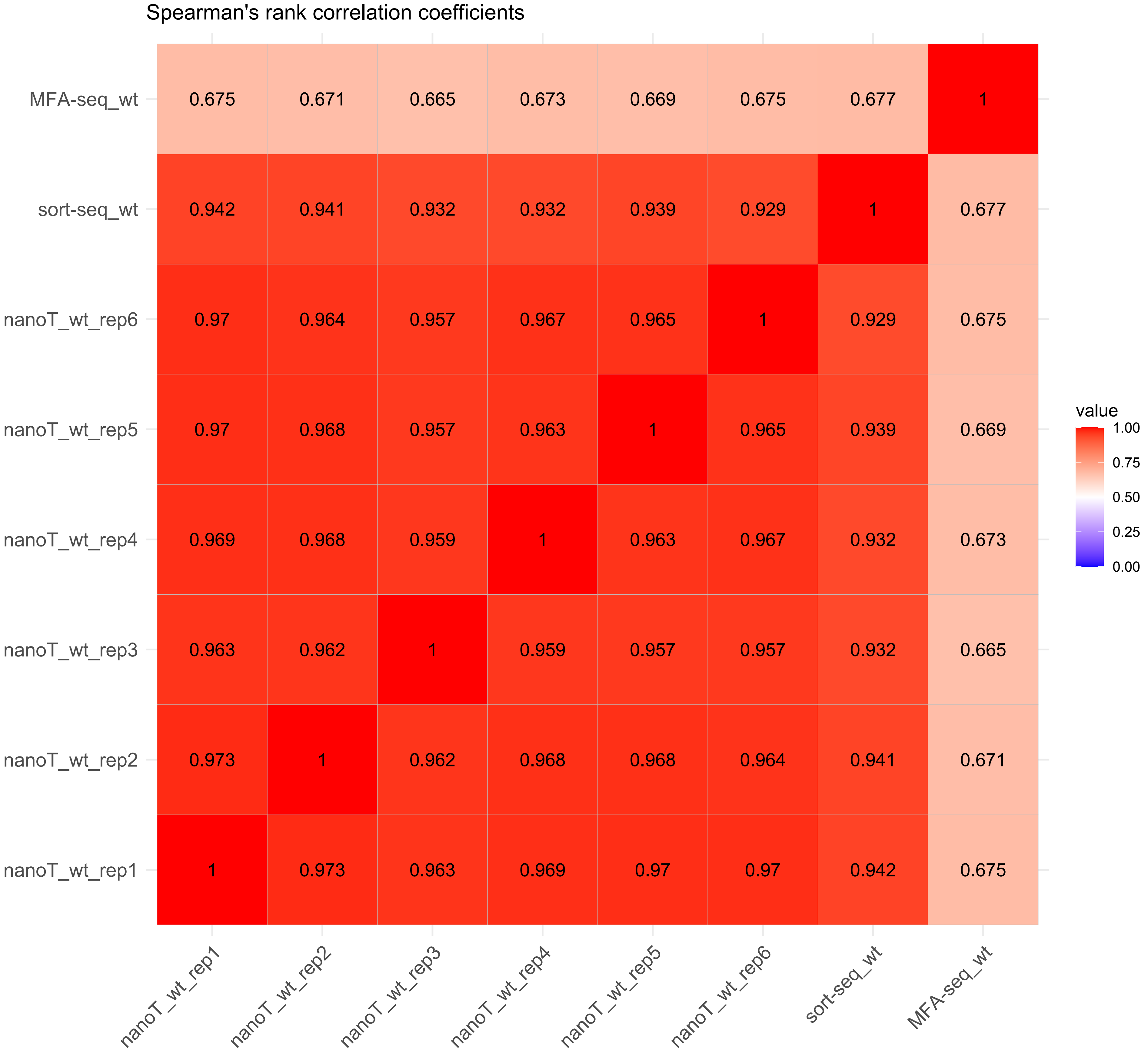

Supplementary Figure 5

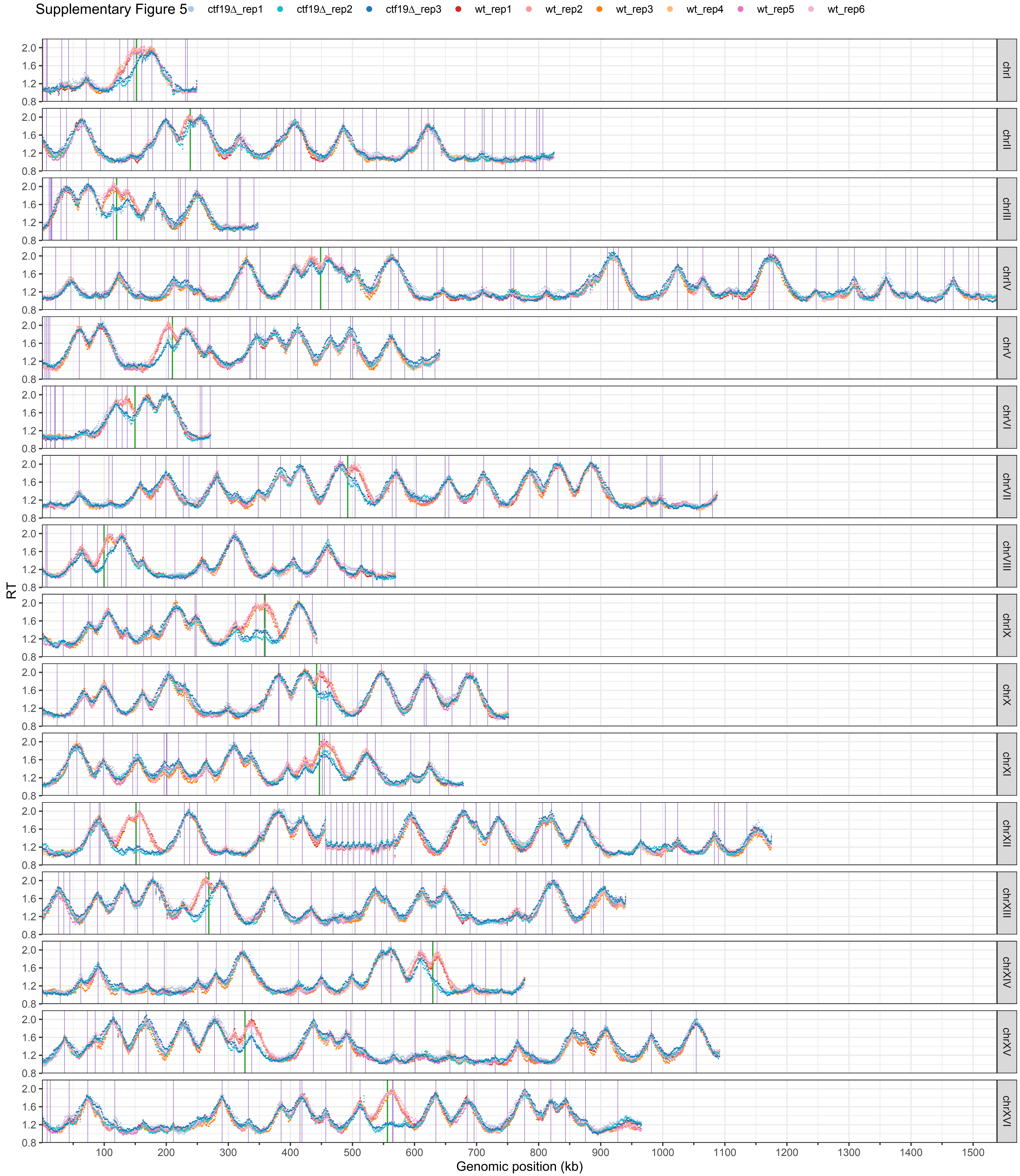

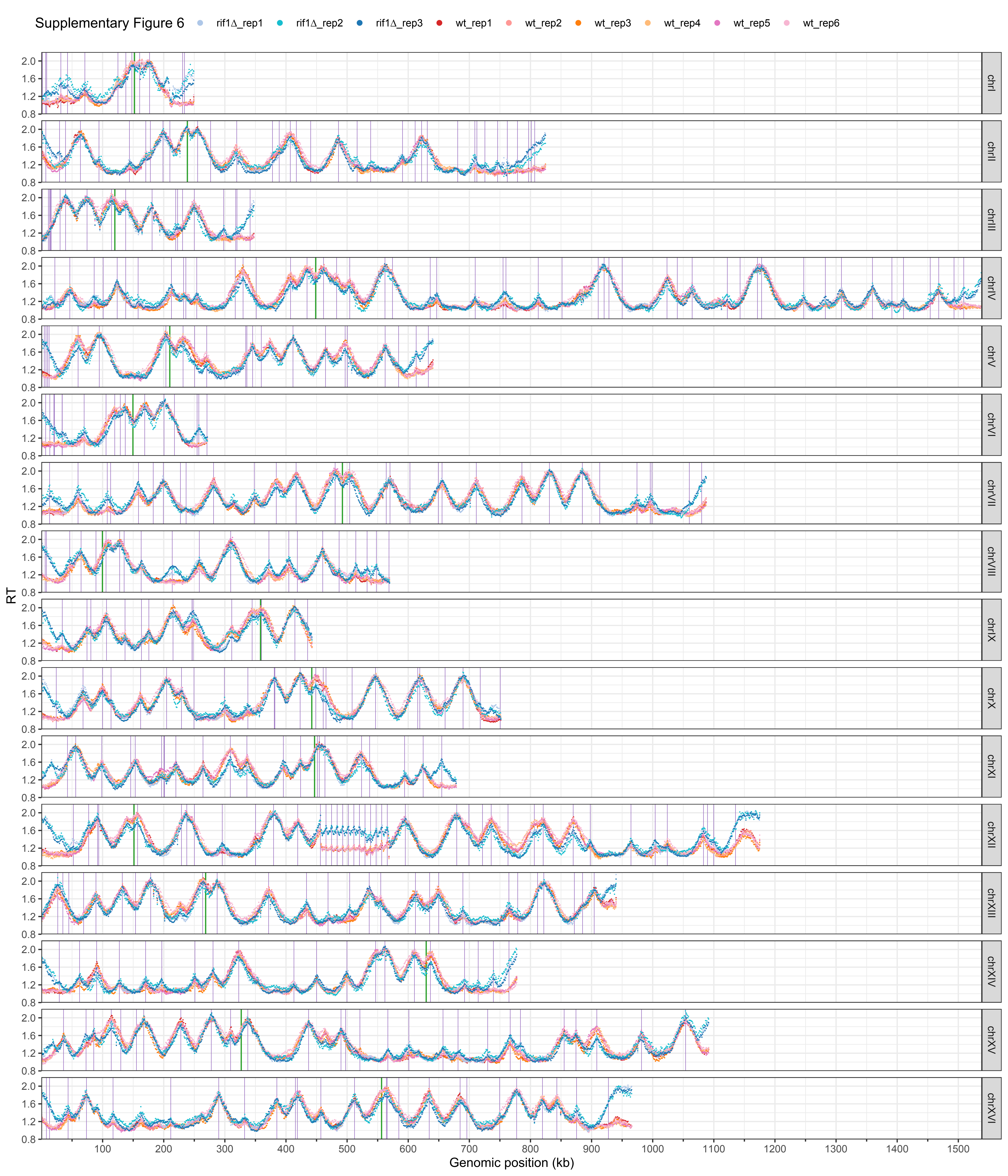
Supplementary Figure 6
rif1Δ_rep1
rif1Δ_rep2
rif1Δ_rep3
wt_rep1
wt_rep2
wt_rep3
wt_rep4
wt_rep5
wt_rep6
RT
Genomic position (kb)
chrI
chrII
chrIII
chrIV
chrV
chrVI
chrVII
chrVIII
chrIX
chrX
chrXI
chrXII
chrXIII
chrXIV
chrXV
chrXVI

Supplementary Figure 7

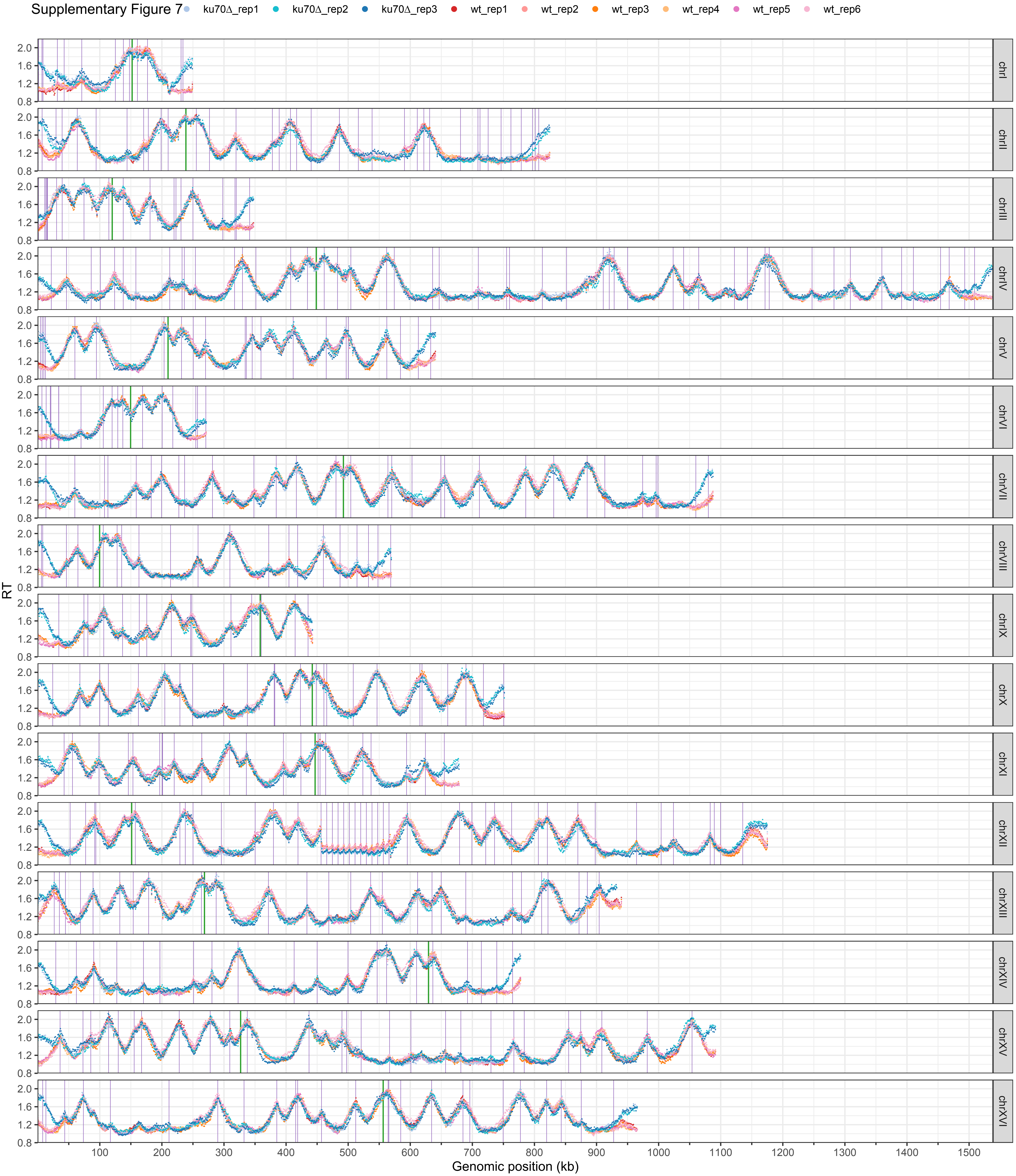

Supplementary Figure 8

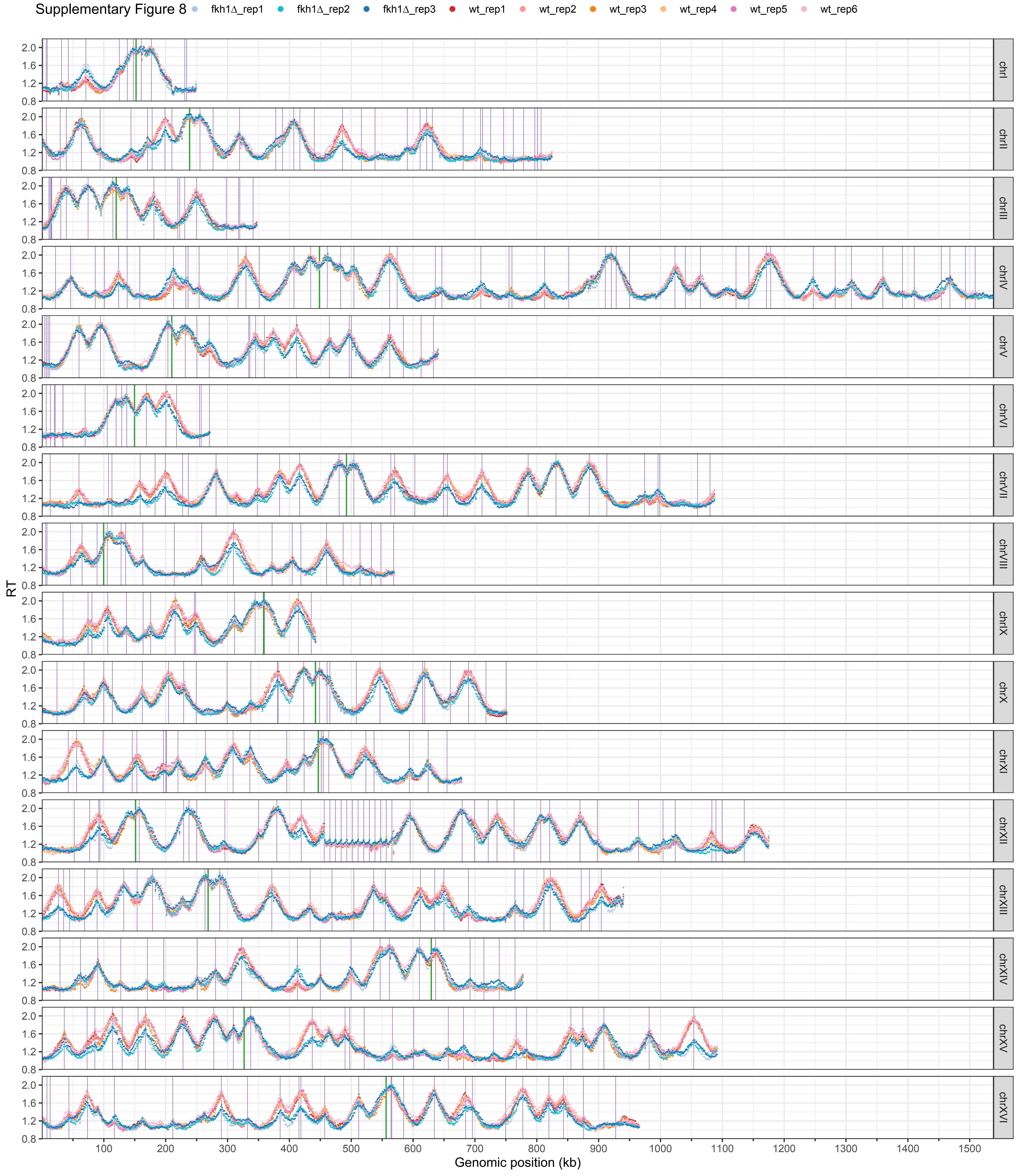

Supplementary Figure 9

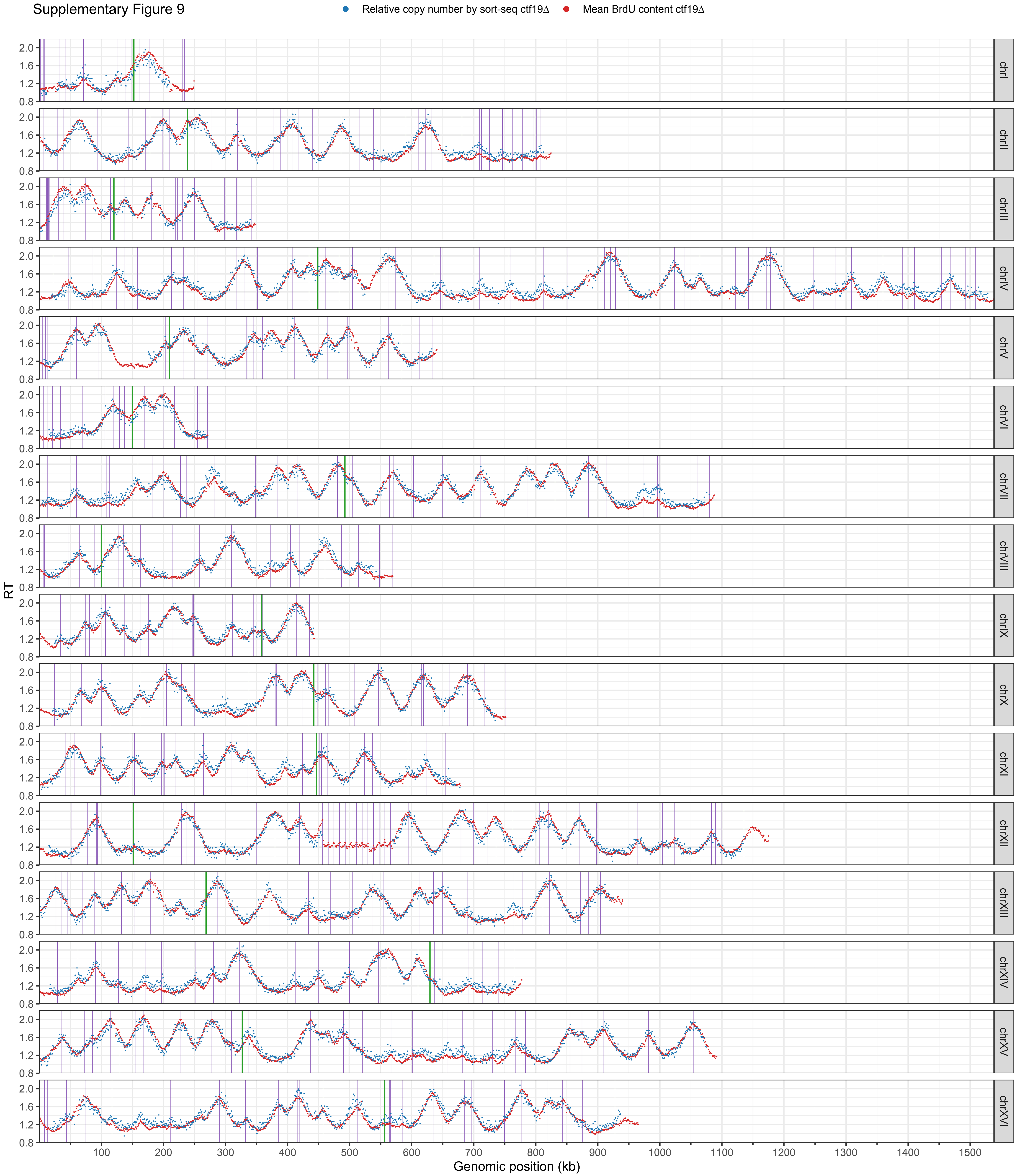

Supplementary Figure 10

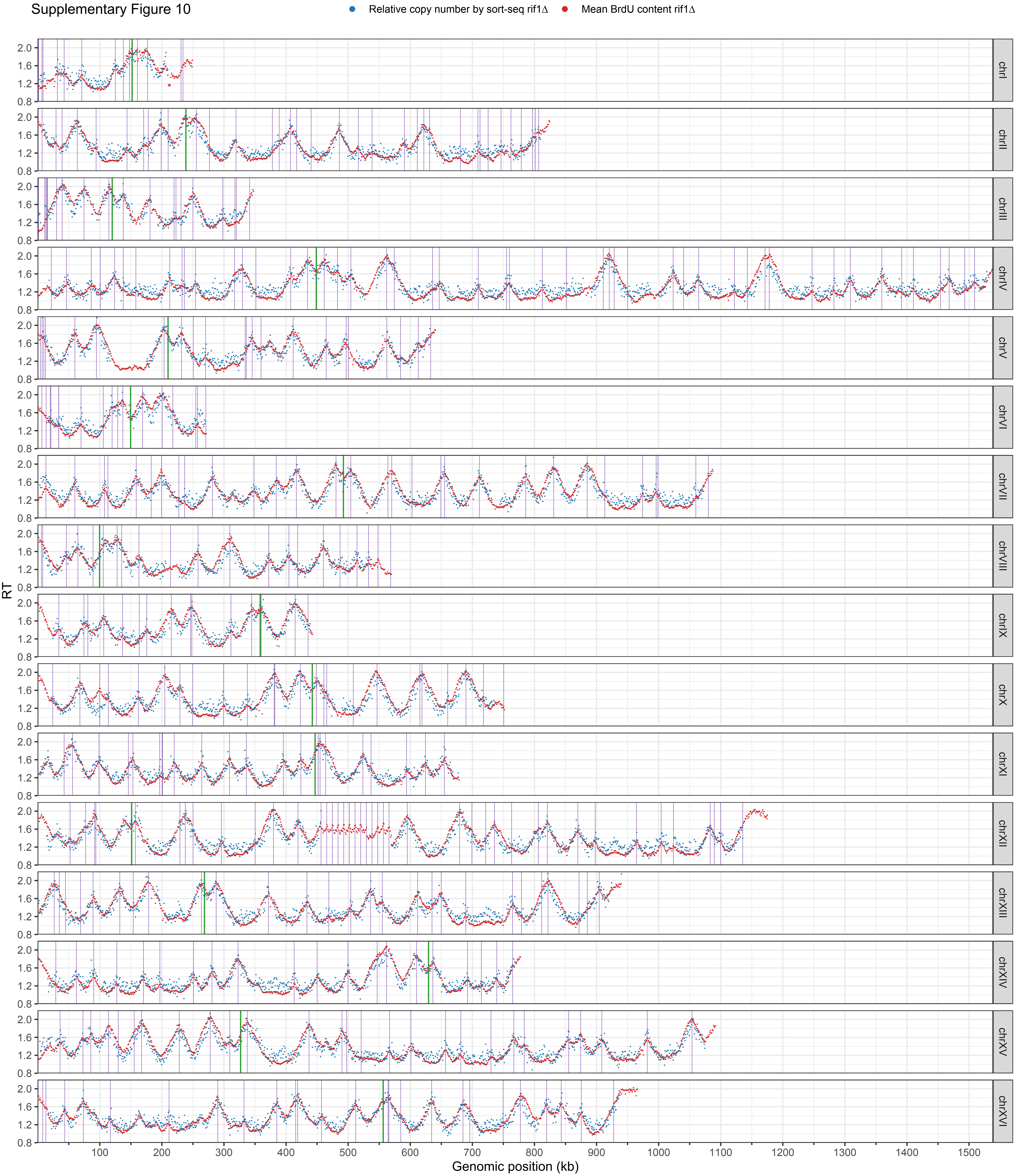

Supplementary Figure 11

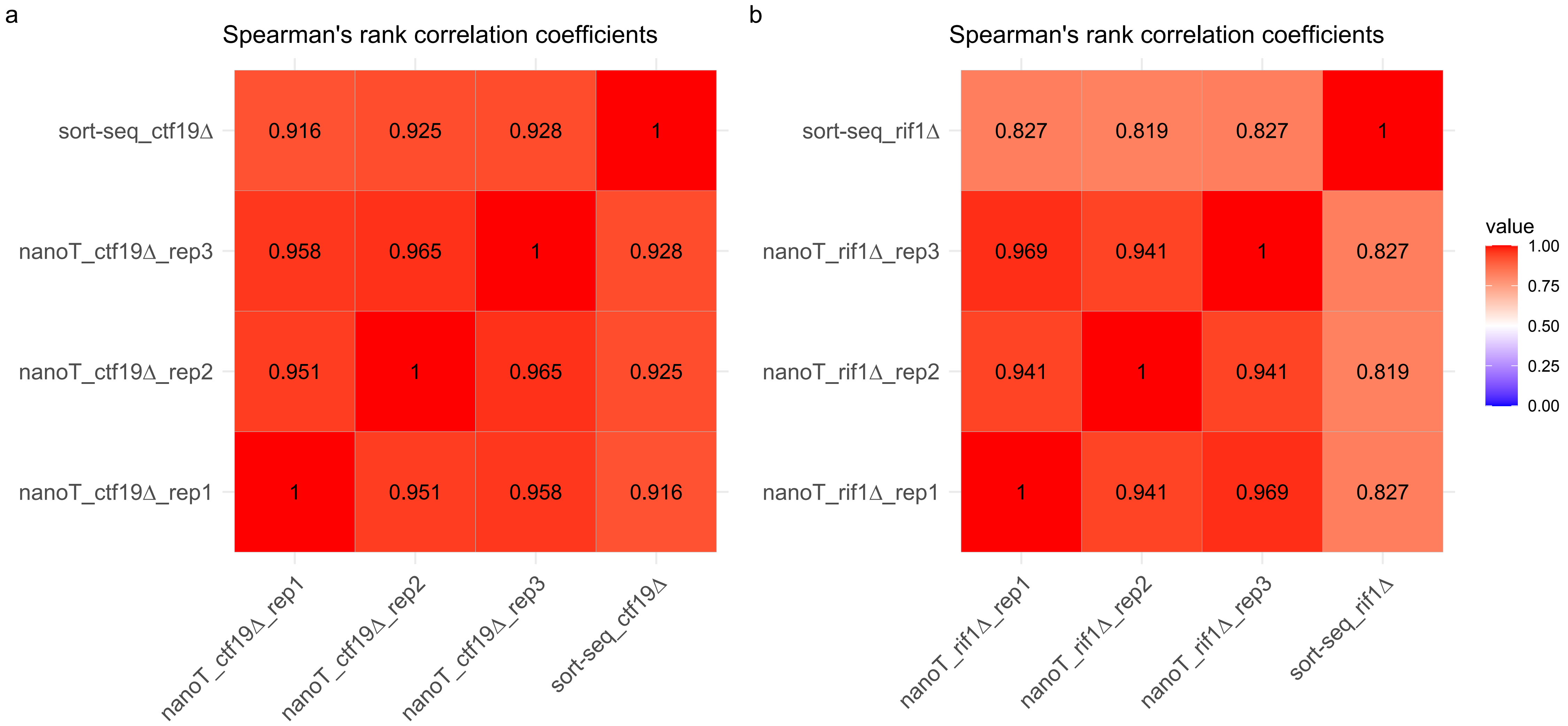

Supplementary Figure 12

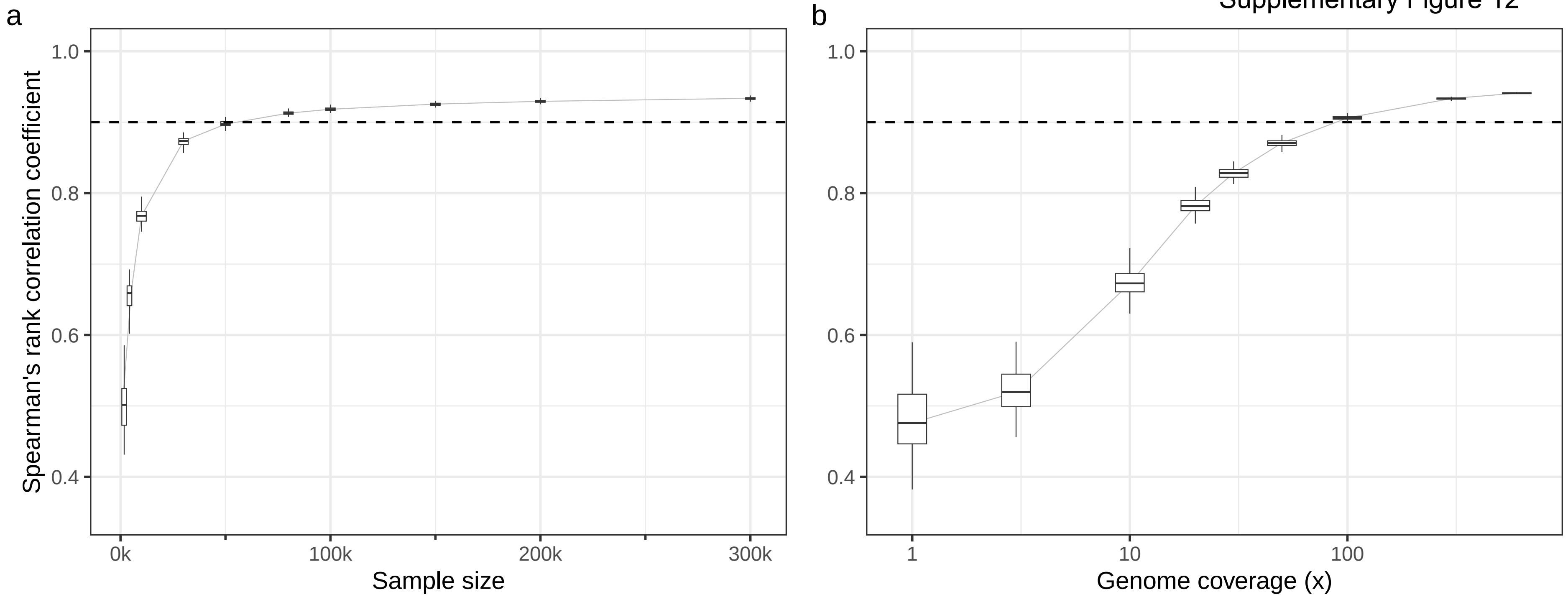

Supplementary Figure 13

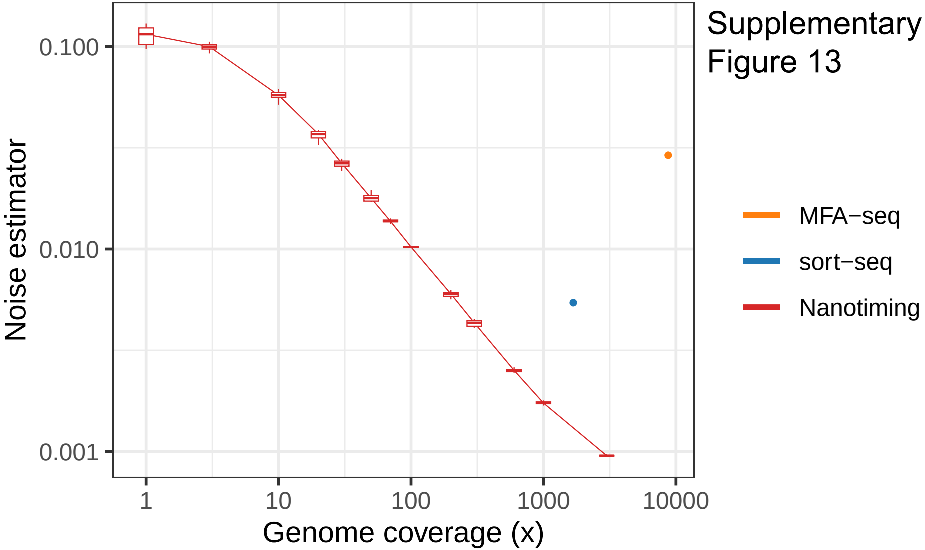

Supplementary Figure 14

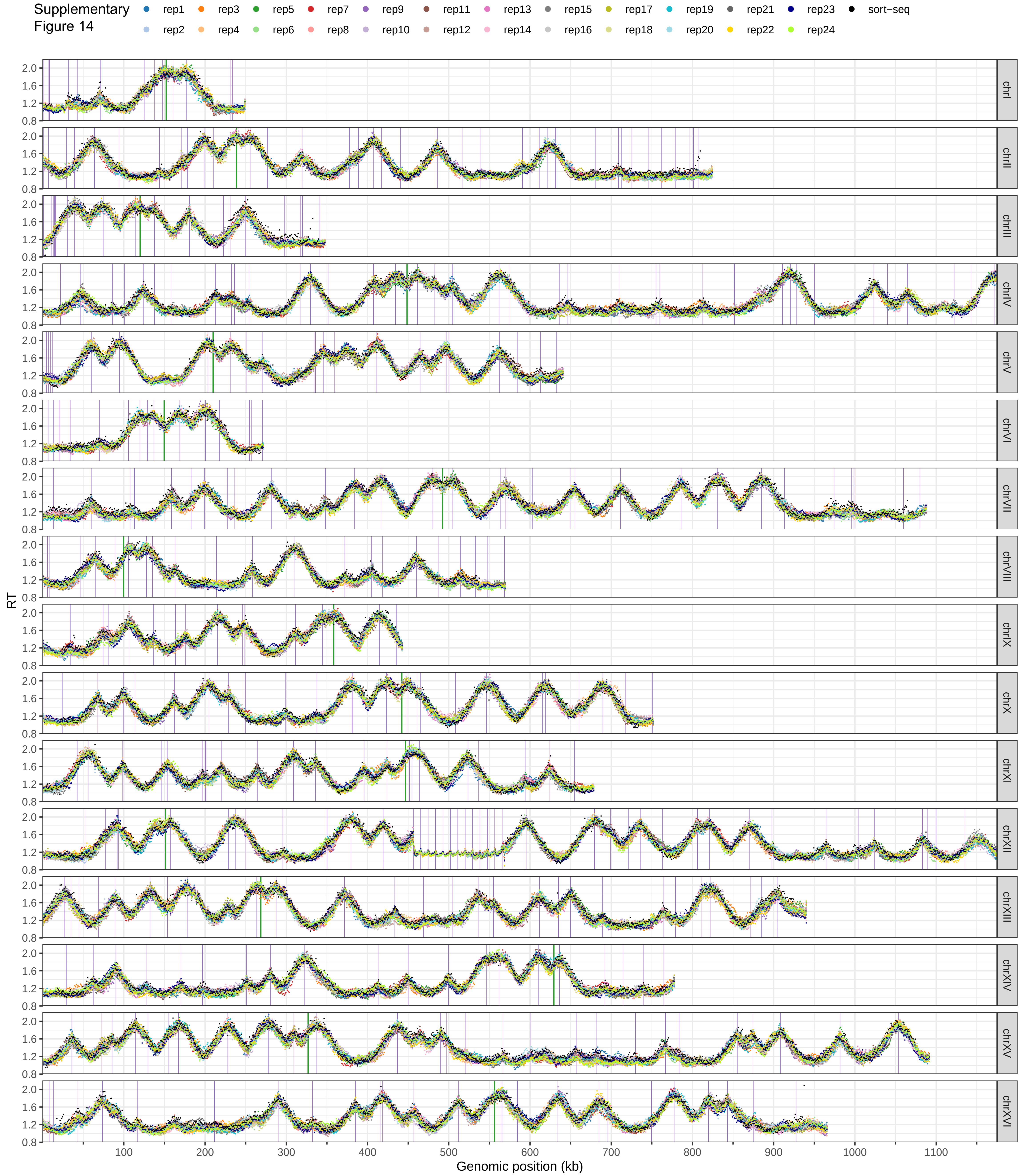

Supplementary Figure 15

Spearman's rank correlation coefficients

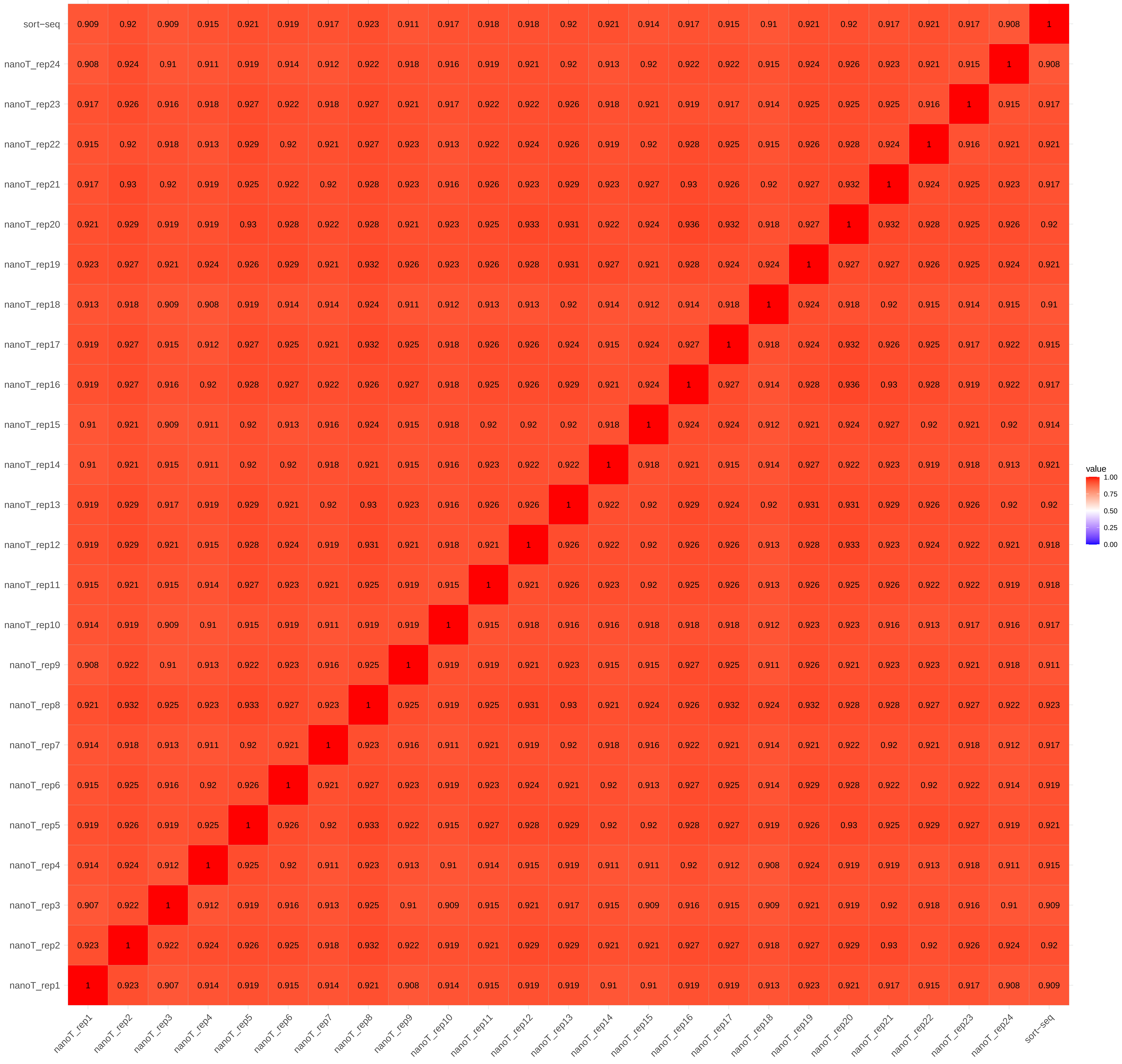

Supplementary Figure 16

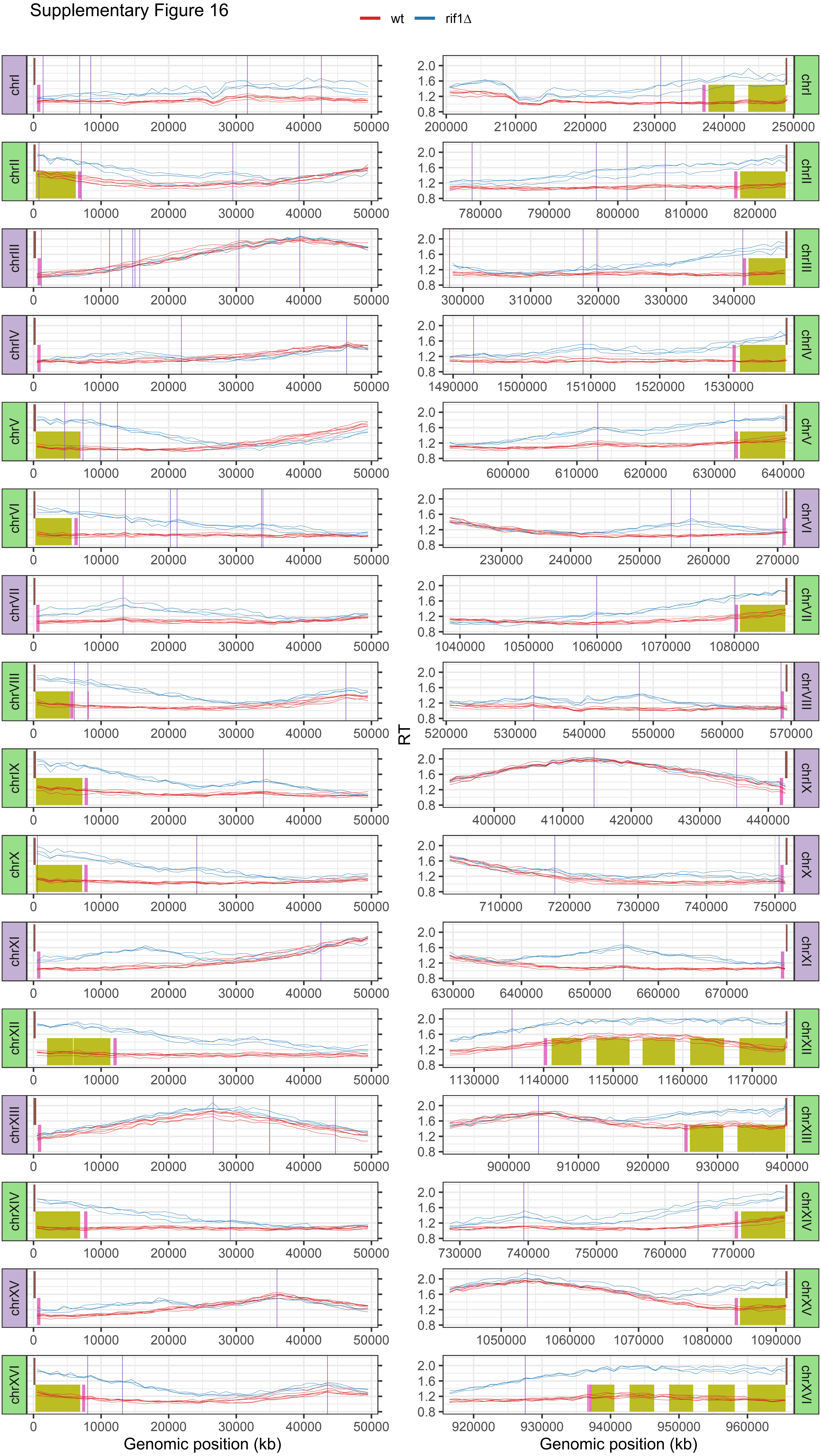

Supplementary Figure 17

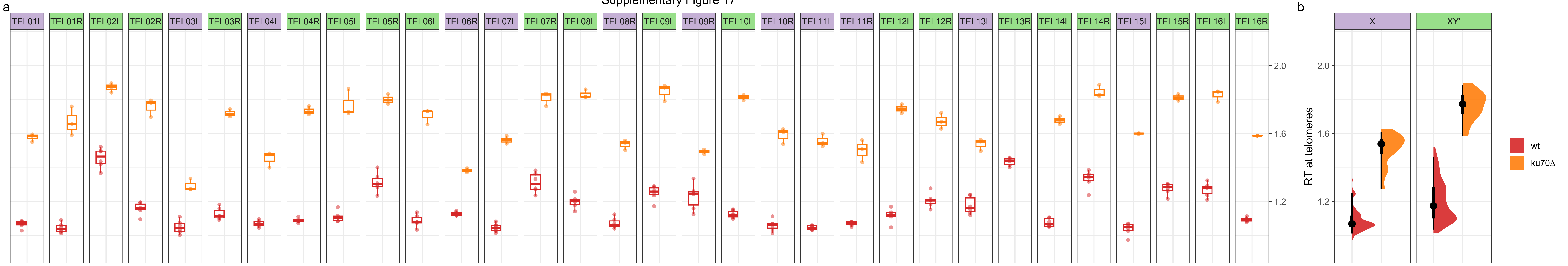

Supplementary Figure 18

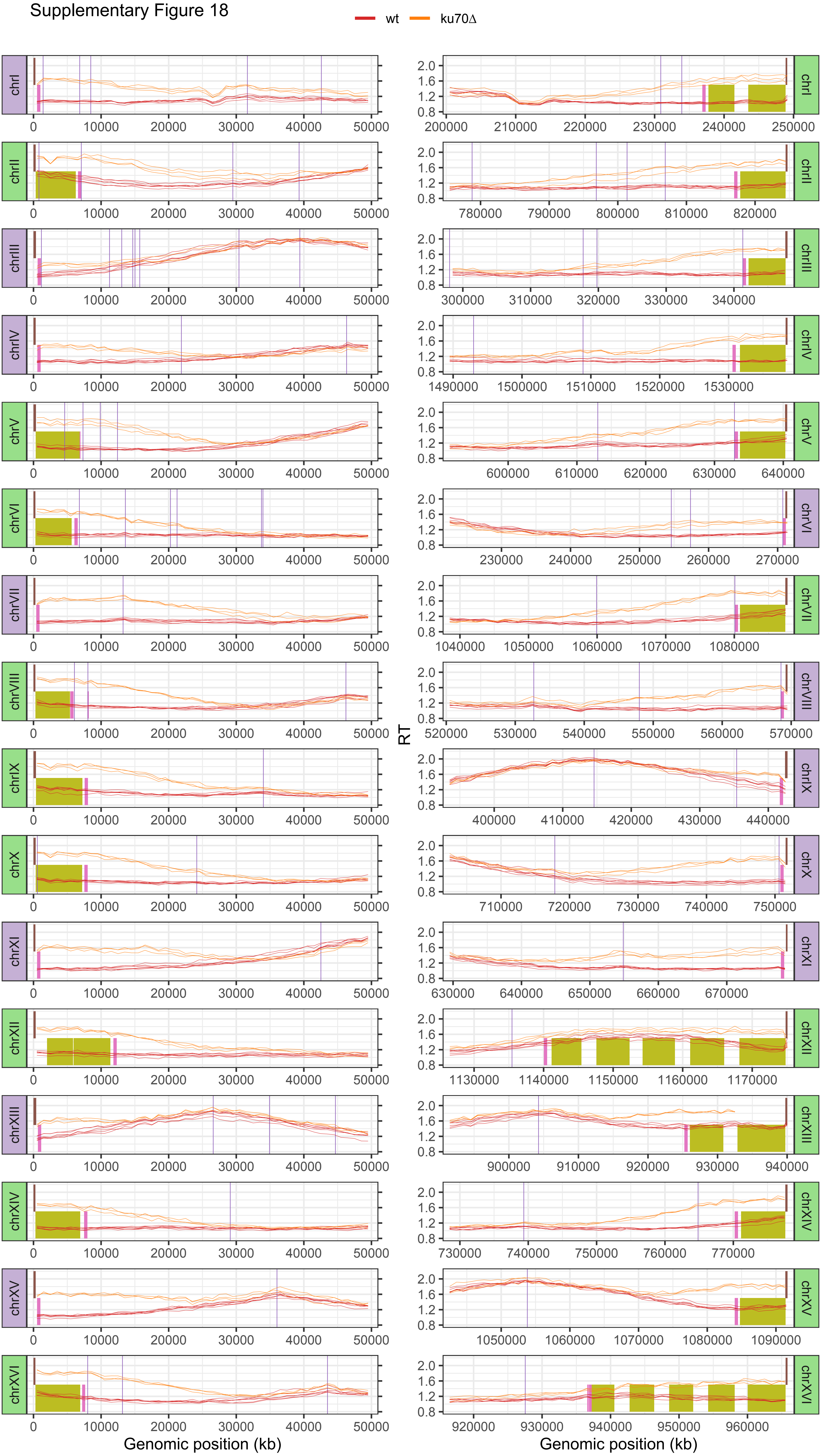

S19: wt

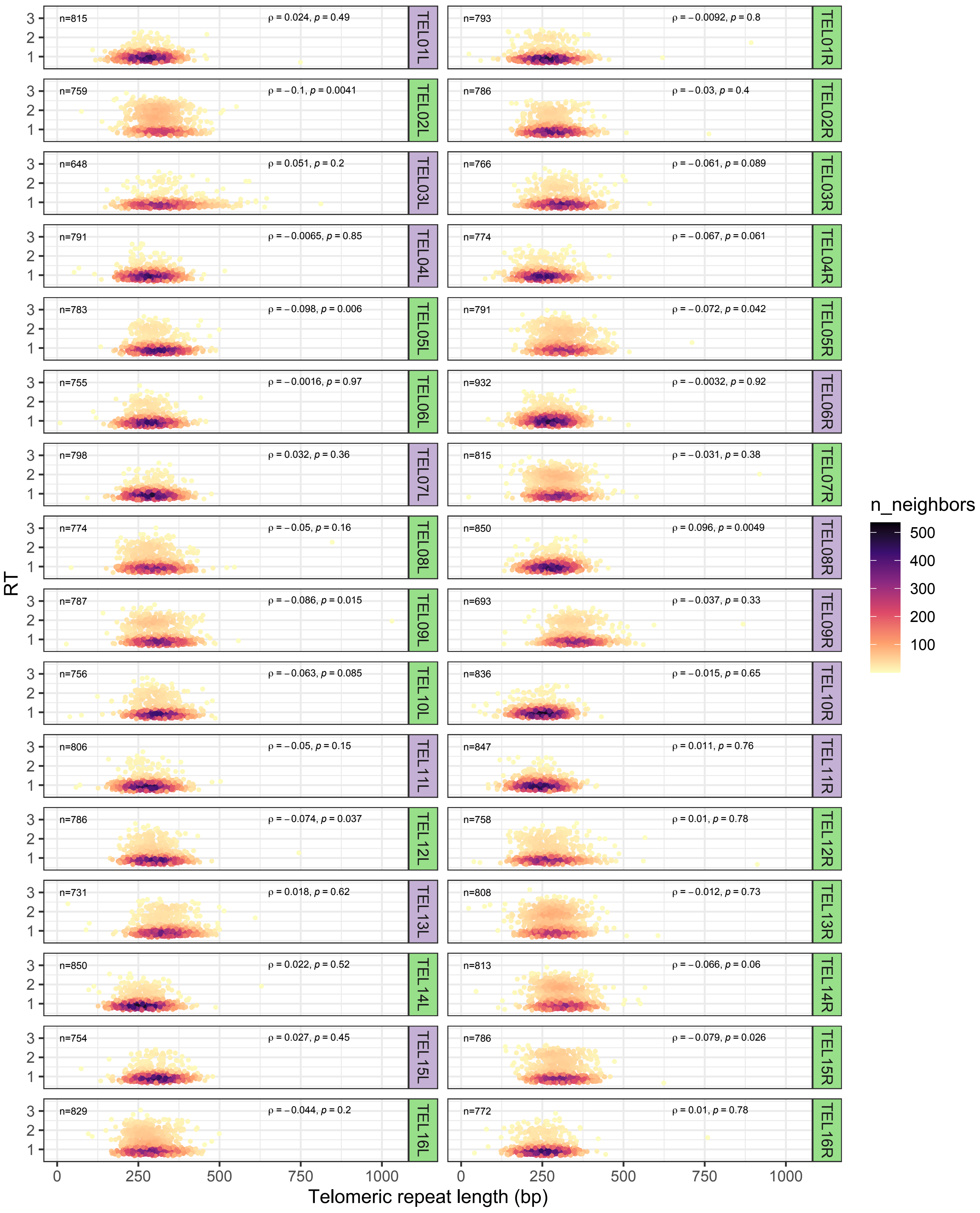

n=815 ρ = 0.024, p = 0.49 TEL01L
n=793 ρ = −0.0092, p = 0.8 TEL01R
n=759 ρ = −0.1, p = 0.0041 TEL02L
n=786 ρ = −0.03, p = 0.4 TEL02R
n=648 ρ = 0.051, p = 0.2 TEL03L
n=766 ρ = −0.061, p = 0.089 TEL03R
n=791 ρ = −0.0065, p = 0.85 TEL04L
n=774 ρ = −0.067, p = 0.061 TEL04R
n=783 ρ = −0.098, p = 0.006 TEL05L
n=791 ρ = −0.072, p = 0.042 TEL05R
n=755 ρ = −0.0016, p = 0.97 TEL06L
n=932 ρ = −0.0032, p = 0.92 TEL06R
n=798 ρ = 0.032, p = 0.36 TEL07L
n=815 ρ = −0.031, p = 0.38 TEL07R
n=774 ρ = −0.05, p = 0.16 TEL08L
n=850 ρ = 0.096, p = 0.0049 TEL08R
n=787 ρ = −0.086, p = 0.015 TEL09L
n=693 ρ = −0.037, p = 0.33 TEL09R
n=756 ρ = −0.063, p = 0.085 TEL10L
n=836 ρ = −0.015, p = 0.65 TEL10R
n=806 ρ = −0.05, p = 0.15 TEL11L
n=847 ρ = 0.011, p = 0.76 TEL11R
n=786 ρ = −0.074, p = 0.037 TEL12L
n=758 ρ = 0.01, p = 0.78 TEL12R
n=731 ρ = 0.018, p = 0.62 TEL13L
n=808 ρ = −0.012, p = 0.73 TEL13R
n=850 ρ = 0.022, p = 0.52 TEL14L
n=813 ρ = −0.066, p = 0.06 TEL14R
n=754 ρ = 0.027, p = 0.45 TEL15L
n=786 ρ = −0.079, p = 0.026 TEL15R
n=829 ρ = −0.044, p = 0.2 TEL16L
n=772 ρ = 0.01, p = 0.78 TEL16R
RT
1
2
3
0
250
500
750
1000
Telomeric repeat length (bp)
n_neighbors
500
400
300
200
100

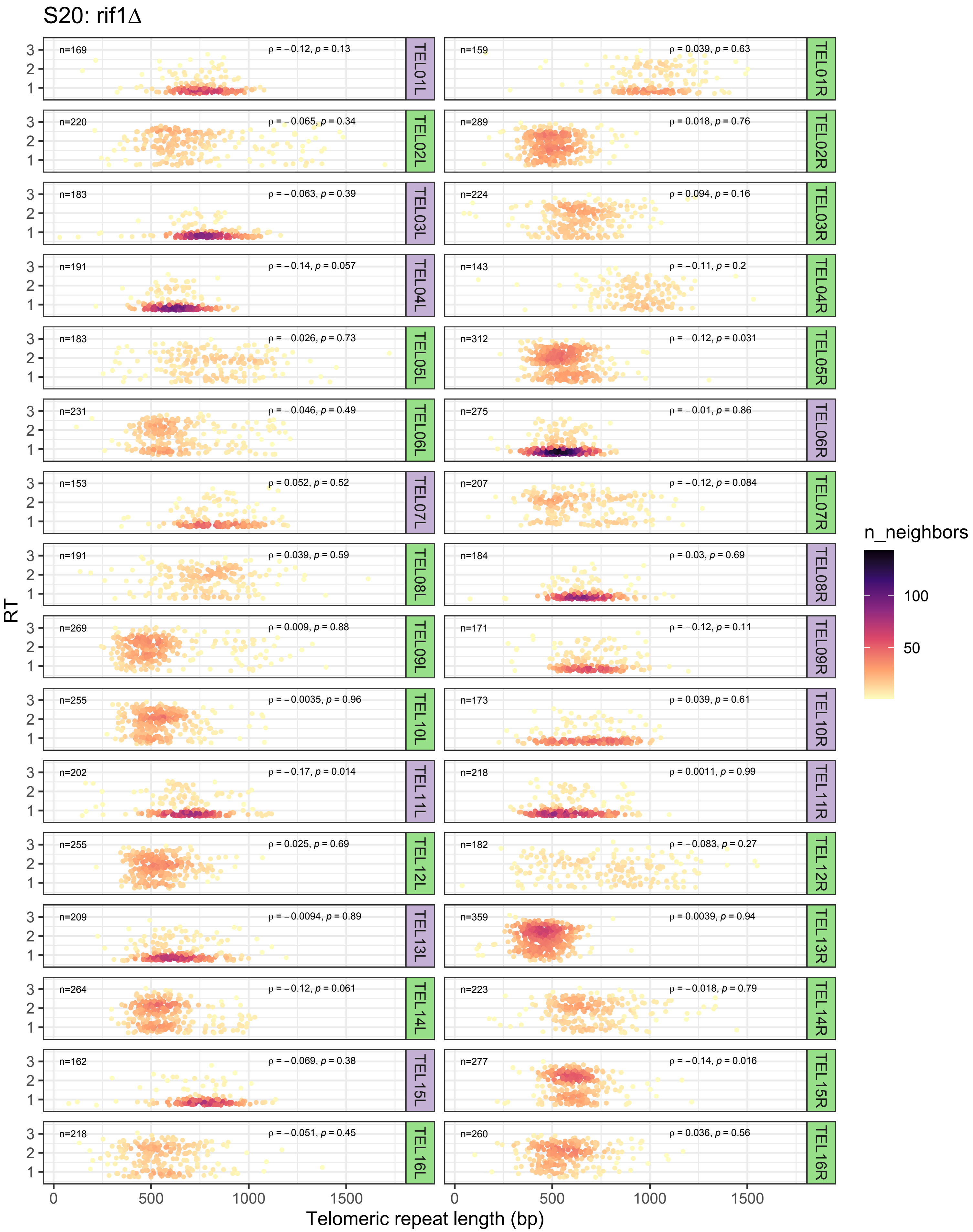

S20: rif1Δ
n=169
ρ = −0.12, p = 0.13
TEL01L
n=159
ρ = 0.039, p = 0.63
TEL01R
n=220
ρ = −0.065, p = 0.34
TEL02L
n=289
ρ = 0.018, p = 0.76
TEL02R
n=183
ρ = −0.063, p = 0.39
TEL03L
n=224
ρ = 0.094, p = 0.16
TEL03R
n=191
ρ = −0.14, p = 0.057
TEL04L
n=143
ρ = −0.11, p = 0.2
TEL04R
n=183
ρ = −0.026, p = 0.73
TEL05L
n=312
ρ = −0.12, p = 0.031
TEL05R
n=231
ρ = −0.046, p = 0.49
TEL06L
n=275
ρ = −0.01, p = 0.86
TEL06R
n=153
ρ = 0.052, p = 0.52
TEL07L
n=207
ρ = −0.12, p = 0.084
TEL07R
n=191
ρ = 0.039, p = 0.59
TEL08L
n=184
ρ = 0.03, p = 0.69
TEL08R
n=269
ρ = 0.009, p = 0.88
TEL09L
n=171
ρ = −0.12, p = 0.11
TEL09R
n=255
ρ = −0.0035, p = 0.96
TEL10L
n=173
ρ = 0.039, p = 0.61
TEL10R
n=202
ρ = −0.17, p = 0.014
TEL11L
n=218
ρ = 0.0011, p = 0.99
TEL11R
n=255
ρ = 0.025, p = 0.69
TEL12L
n=182
ρ = −0.083, p = 0.27
TEL12R
n=209
ρ = −0.0094, p = 0.89
TEL13L
n=359
ρ = 0.0039, p = 0.94
TEL13R
n=264
ρ = −0.12, p = 0.061
TEL14L
n=223
ρ = −0.018, p = 0.79
TEL14R
n=162
ρ = −0.069, p = 0.38
TEL15L
n=277
ρ = −0.14, p = 0.016
TEL15R
n=218
ρ = −0.051, p = 0.45
TEL16L
n=260
ρ = 0.036, p = 0.56
TEL16R
RT
1
2
3
0
500
1000
1500
Telomeric repeat length (bp)
n_neighbors
100
50

S21: ku70Δ

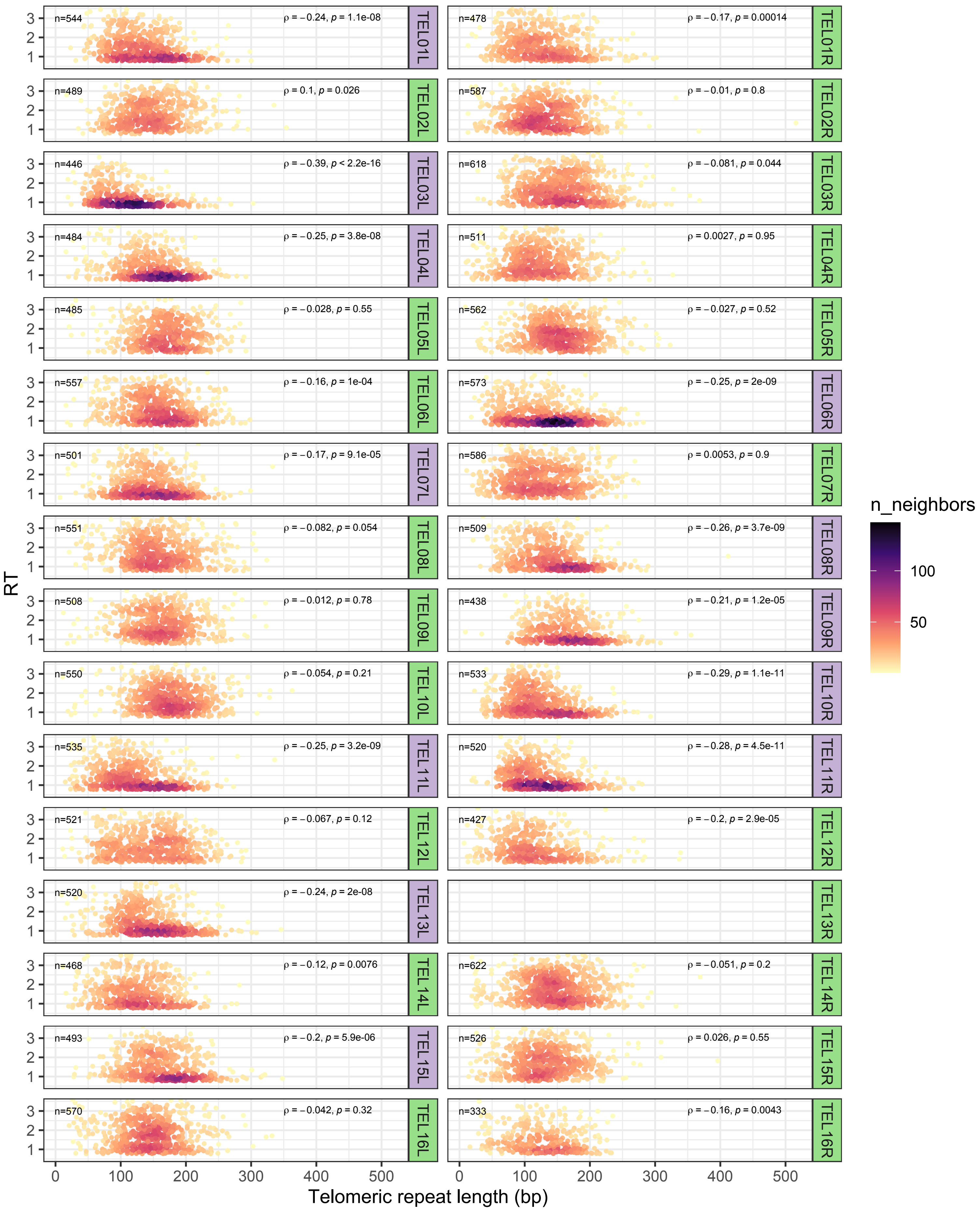

n=544
ρ = −0.24, p = 1.1e-08
TEL01L
n=478
ρ = −0.17, p = 0.00014
TEL01R
n=489
ρ = 0.1, p = 0.026
TEL02L
n=587
ρ = −0.01, p = 0.8
TEL02R
n=446
ρ = −0.39, p < 2.2e-16
TEL03L
n=618
ρ = −0.081, p = 0.044
TEL03R
n=484
ρ = −0.25, p = 3.8e-08
TEL04L
n=511
ρ = 0.0027, p = 0.95
TEL04R
n=485
ρ = −0.028, p = 0.55
TEL05L
n=562
ρ = −0.027, p = 0.52
TEL05R
n=557
ρ = −0.16, p = 1e-04
TEL06L
n=573
ρ = −0.25, p = 2e-09
TEL06R
n=501
ρ = −0.17, p = 9.1e-05
TEL07L
n=586
ρ = 0.0053, p = 0.9
TEL07R
n=551
ρ = −0.082, p = 0.054
TEL08L
n=509
ρ = −0.26, p = 3.7e-09
TEL08R
n=508
ρ = −0.012, p = 0.78
TEL09L
n=438
ρ = −0.21, p = 1.2e-05
TEL09R
n=550
ρ = −0.054, p = 0.21
TEL10L
n=533
ρ = −0.29, p = 1.1e-11
TEL10R
n=535
ρ = −0.25, p = 3.2e-09
TEL11L
n=520
ρ = −0.28, p = 4.5e-11
TEL11R
n=521
ρ = −0.067, p = 0.12
TEL12L
n=427
ρ = −0.2, p = 2.9e-05
TEL12R
n=520
ρ = −0.24, p = 2e-08
TEL13L
TEL13R
n=468
ρ = −0.12, p = 0.0076
TEL14L
n=622
ρ = −0.051, p = 0.2
TEL14R
n=493
ρ = −0.2, p = 5.9e-06
TEL15L
n=526
ρ = 0.026, p = 0.55
TEL15R
n=570
ρ = −0.042, p = 0.32
TEL16L
n=333
ρ = −0.16, p = 0.0043
TEL16R
RT
1
2
3
0
100
200
300
400
500
Telomeric repeat length (bp)
n_neighbors
100
50

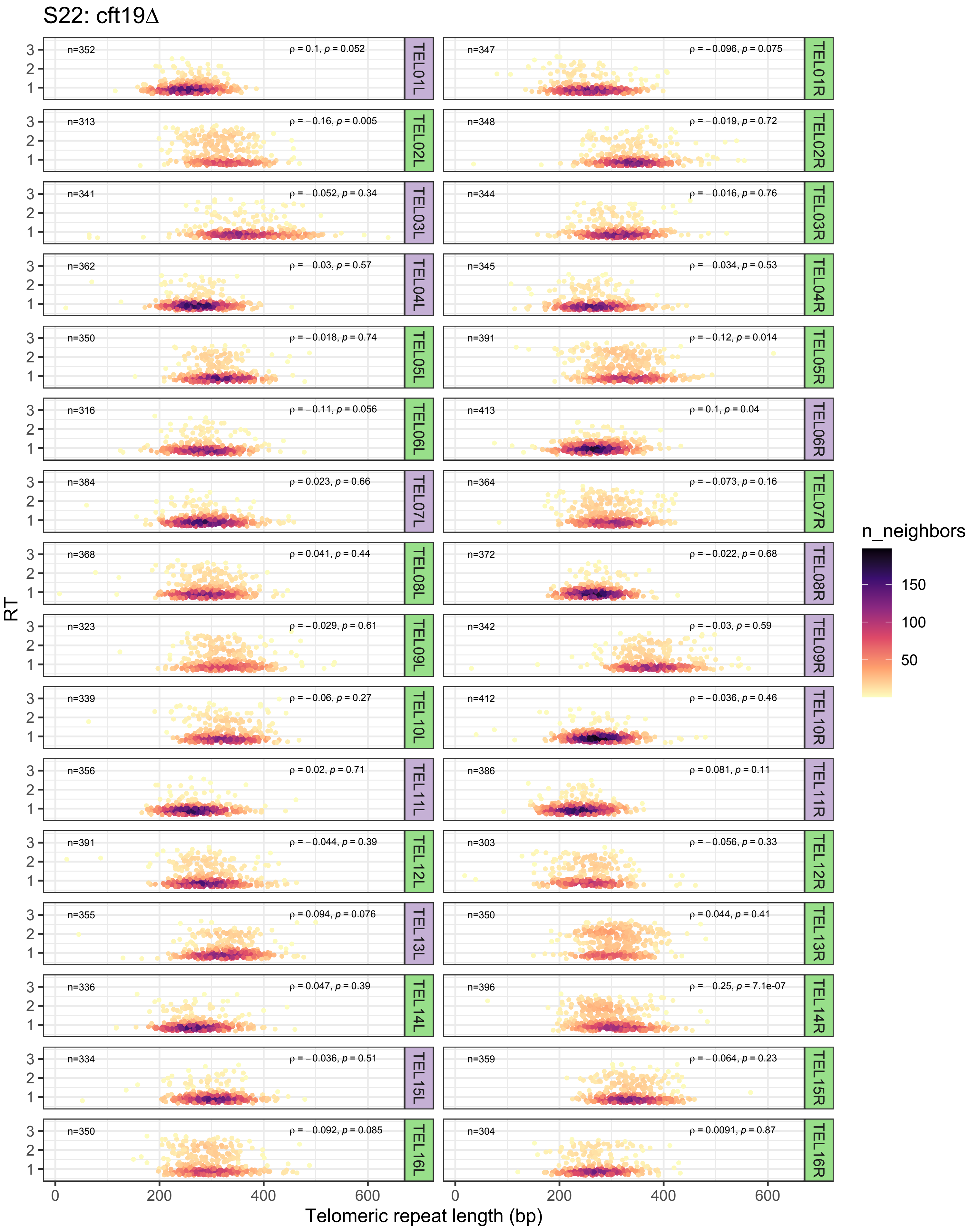

S22: cft19Δ
n=352
ρ = 0.1, p = 0.052
TEL01L
n=347
ρ = −0.096, p = 0.075
TEL01R
n=313
ρ = −0.16, p = 0.005
TEL02L
n=348
ρ = −0.019, p = 0.72
TEL02R
n=341
ρ = −0.052, p = 0.34
TEL03L
n=344
ρ = −0.016, p = 0.76
TEL03R
n=362
ρ = −0.03, p = 0.57
TEL04L
n=345
ρ = −0.034, p = 0.53
TEL04R
n=350
ρ = −0.018, p = 0.74
TEL05L
n=391
ρ = −0.12, p = 0.014
TEL05R
n=316
ρ = −0.11, p = 0.056
TEL06L
n=413
ρ = 0.1, p = 0.04
TEL06R
n=384
ρ = 0.023, p = 0.66
TEL07L
n=364
ρ = −0.073, p = 0.16
TEL07R
n=368
ρ = 0.041, p = 0.44
TEL08L
n=372
ρ = −0.022, p = 0.68
TEL08R
n=323
ρ = −0.029, p = 0.61
TEL09L
n=342
ρ = −0.03, p = 0.59
TEL09R
n=339
ρ = −0.06, p = 0.27
TEL10L
n=412
ρ = −0.036, p = 0.46
TEL10R
n=356
ρ = 0.02, p = 0.71
TEL11L
n=386
ρ = 0.081, p = 0.11
TEL11R
n=391
ρ = −0.044, p = 0.39
TEL12L
n=303
ρ = −0.056, p = 0.33
TEL12R
n=355
ρ = 0.094, p = 0.076
TEL13L
n=350
ρ = 0.044, p = 0.41
TEL13R
n=336
ρ = 0.047, p = 0.39
TEL14L
n=396
ρ = −0.25, p = 7.1e-07
TEL14R
n=334
ρ = −0.036, p = 0.51
TEL15L
n=359
ρ = −0.064, p = 0.23
TEL15R
n=350
ρ = −0.092, p = 0.085
TEL16L
n=304
ρ = 0.0091, p = 0.87
TEL16R
RT
1
2
3
0
200
400
600
Telomeric repeat length (bp)
n_neighbors
150
100
50

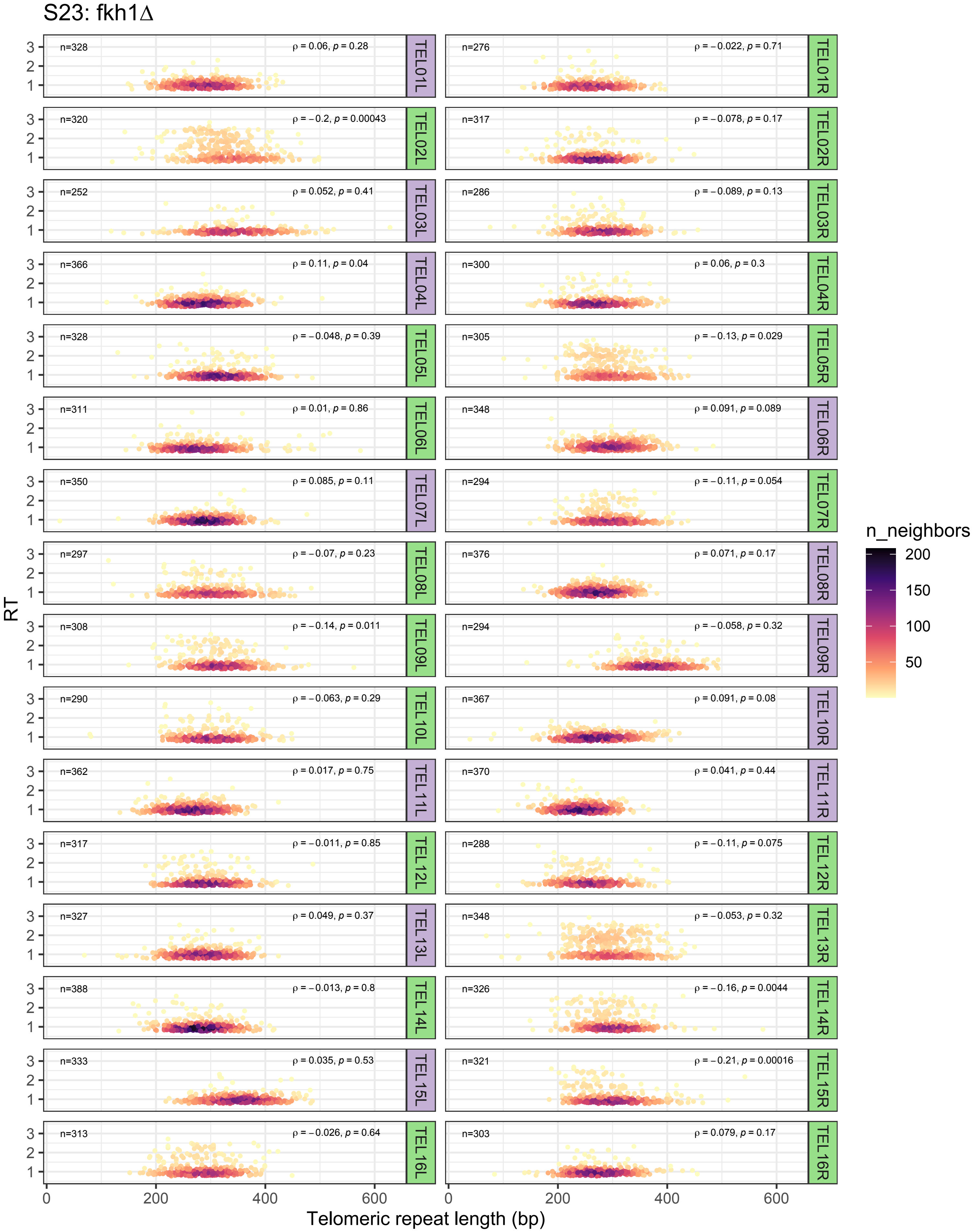
S23: fkh1Δ
RT
Telomeric repeat length (bp)
n_neighbors
TEL01L n=328 ρ = 0.06, p = 0.28
TEL01R n=276 ρ = −0.022, p = 0.71
TEL02L n=320 ρ = −0.2, p = 0.00043
TEL02R n=317 ρ = −0.078, p = 0.17
TEL03L n=252 ρ = 0.052, p = 0.41
TEL03R n=286 ρ = −0.089, p = 0.13
TEL04L n=366 ρ = 0.11, p = 0.04
TEL04R n=300 ρ = 0.06, p = 0.3
TEL05L n=328 ρ = −0.048, p = 0.39
TEL05R n=305 ρ = −0.13, p = 0.029
TEL06L n=311 ρ = 0.01, p = 0.86
TEL06R n=348 ρ = 0.091, p = 0.089
TEL07L n=350 ρ = 0.085, p = 0.11
TEL07R n=294 ρ = −0.11, p = 0.054
TEL08L n=297 ρ = −0.07, p = 0.23
TEL08R n=376 ρ = 0.071, p = 0.17
TEL09L n=308 ρ = −0.14, p = 0.011
TEL09R n=294 ρ = −0.058, p = 0.32
TEL10L n=290 ρ = −0.063, p = 0.29
TEL10R n=367 ρ = 0.091, p = 0.08
TEL11L n=362 ρ = 0.017, p = 0.75
TEL11R n=370 ρ = 0.041, p = 0.44
TEL12L n=317 ρ = −0.011, p = 0.85
TEL12R n=288 ρ = −0.11, p = 0.075
TEL13L n=327 ρ = 0.049, p = 0.37
TEL13R n=348 ρ = −0.053, p = 0.32
TEL14L n=388 ρ = −0.013, p = 0.8
TEL14R n=326 ρ = −0.16, p = 0.0044
TEL15L n=333 ρ = 0.035, p = 0.53
TEL15R n=321 ρ = −0.21, p = 0.00016
TEL16L n=313 ρ = −0.026, p = 0.64
TEL16R n=303 ρ = 0.079, p = 0.17

### Supplementary Figure 24

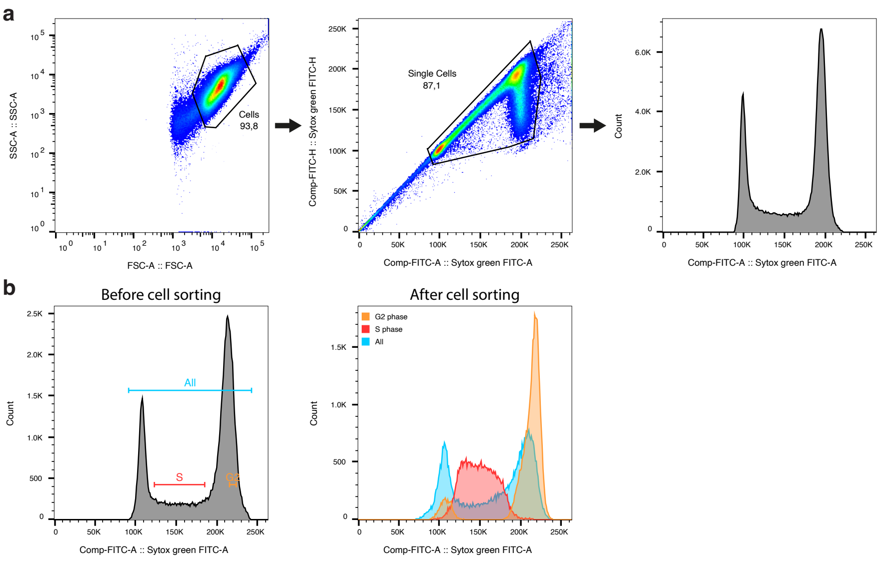
